# Regulation of lateral root development by shoot-sensed far-red light via HY5 is nitrate-dependent and involves the NRT2.1 nitrate transporter

**DOI:** 10.1101/2021.01.31.428985

**Authors:** Kasper van Gelderen, Chiakai Kang, Peijin Li, Ronald Pierik

## Abstract

Plants are very effective in responding to environmental changes during competition for light and nutrients. Low Red:Far-Red (low R:FR)-mediated neighbor detection allows plants to compete successfully with other plants for available light. This above-ground signal can also reduce lateral root growth by inhibiting lateral root emergence, a process that might help the plant invest resources in shoot growth. Nitrate is an essential nutrient for plant growth and *Arabidopsis thaliana* responds to low nitrate conditions by enhancing nutrient uptake and reducing lateral and main root growth. There are indications that low R:FR signaling and low nitrate signaling can affect each other. It is unknown which response is prioritized when low R:FR light- and low nitrate signaling co-occur. We investigated the effect of low nitrate conditions on the low R:FR response of the *A. thaliana* root system in agar plate media, combined with the application of supplemental Far-Red (FR) light to the shoot. We observed that under low nitrate conditions main and lateral root growth was reduced, but more importantly, that the response of the root system to low R:FR was suppressed. Consistently, a loss-of-function mutant of a nitrate transporter gene *NRT2.1* lacked low R:FR-induced lateral root reduction and its root growth was hypersensitive to low nitrate. ELONGATED HYPOCOTYL5 (HY5) plays an important role in the root response to low R:FR and we found that it was less sensitive to low nitrate conditions with regards to lateral root growth. In addition, we found that low R:FR increases *NRT2.1* expression and that low nitrate enhances *HY5* expression. HY5 also affects *NRT2.1* expression, however, it depended on the presence of ammonium in which direction this effect was. Replacing part of the nitrogen source with ammonium also removed the effect of low R:FR on the root system, showing that changes in nitrogen sources can be crucial for root plasticity. Together our results show that nitrate signaling can repress low R:FR responses and that this involves signaling via HY5 and NRT2.1.

## 1 Introduction

Plants adapt their growth and development to compete for the limited light and nutrients with which they grow their bodies. Plant can sense competing neighbours via Far-Red (FR) light that is reflected by leaves of neighbouring plants. This reflection of FR light leads to a lowering of the Red to Far-Red ratio (R:FR). Plants respond to this low R:FR by elongating their aboveground organs in an effort to reach for the sunlight. This adaptive response to future competition is what we call the shade avoidance response (Ballaré and Pierik, 2017). The R:FR ratio is sensed by Phytochrome photoreceptors; Phytochromes are activated by R light, changing them to active Pfr state and inactivated by FR light which changes them back to inactive Pr state. The active Pfr form of phytochromes phosphorylates and interacts with PHYTOCHROME INTERACTING FACTORS (PIFs), leading to their mutual degradation (Chen and Chory, 2011). PIFs which are bHLH transcription factors that regulate light and temperature responses (Leivar and Monte, 2014).

Plant roots are essential for the uptake of water and nutrients from the soil, but root growth is impossible without sugars supplied from the shoot. This interdependency between root and shoot means that signaling between these organs is essential to achieve optimal growth (Van Gelderen et al., 2018). The root system responds to low R:FR-mediated plant competition via by reducing its growth (Salisbury et al., 2007; van Gelderen et al., 2018). Normally the root system cannot directly detect the above-ground low R:FR ratio, therefore a mobile, FR-induced, bZip transcription factor ELONGHATED HYPOCOTYL 5 (HY5) travels from shoot to root to affect root growth belowground (Chen et al., 2016; van Gelderen et al., 2018). In the root, HY5 increases its own expression (Zhang et al., 2017) and represses auxin signaling and lateral root development (Sibout et al., 2006; Cluis et al., 2004). The current model is that FR light enhances HY5 transport to the root, which leads to repression of lateral root emergence by repressing auxin signaling and transport around the developing lateral root primordium (LRP) (van Gelderen et al., 2018).

HY5 transport can also affect nutrient uptake by upregulating transcription of the nitrate transporter gene *NRT2.1* (Chen et al., 2016; Jonassen et al., 2008, 2009). Nitrate is a crucial resource for plant life which is taken up by the root and transported through the xylem to the shoot. There are several transmembrane nitrate transporters that facilitate this uptake. NRT1.1 is a transporter/receptor that plays a crucial role in constant high-affinity nitrate uptake, when nitrate is sufficient (Krouk et al., 2006). NRT2.1 is an important high affinity nitrate transporter in the root (Cerezo et al., 2001) that is upregulated when nitrate concentrations are low and NRT2.1 is crucial for low nitrate responses (O’Brien et al., 2016). Another way that shoot-derived HY5 can regulate nutrient uptake is by upregulating the transcription of the phosphate transporter gene *PHT1*, much in the same manner as in the case of NRT2.1 (Sakuraba et al., 2018). Therefore, it is clear that shoot-perceived low R:FR could regulate nutrient uptake via the root through shoot-to-root transport of HY5.Thus, if light quality can influence nutrient uptake-associated transporters, can nutrient signaling affect low R:FR-mediated changes in root development? In order to test this hypothesis, we grew *Arabidopsis thaliana* in the D-root petri-plate system that allows roots to be kept in darkness despite the plant being on an agar plate (Silva-Navas et al., 2015). In this way only the shoot, and not the root, is experiencing a low R:FR ratio, which we achieve by the addition of supplemental FR to the white light background (WL+FR)(van Gelderen et al., 2018). We combined this setup with different nitrate-containing media and observed that low nitrate inhibited the response of the root and shoot to shoot-perceived WL+FR. Through mutant analyses we were able to show that in addition to HY5, NRT2.1 is also involved in WL+FR-mediated root growth reduction. qRT-PCR Expression analysis showed that both WL+FR light and low nitrate induce *NRT2.1* expression. Additionally, low nitrate induced expression of *HY5*, which linked changes in *NRT2.1* and *HY5* expression and lateral root development phenotypes. Interestingly, the role of HY5 in regulating *NRT2.1* expression was highly dependent on the nitrogen source used (ammonium and/or nitrate). Together these results provide a causal link for the integration of WL+FR signaling from the shoot with nutrient signaling in the root via HY5 and NRT2.1.

## 2 Materials and Methods

### 2.1 Plant material

In all experiments Columbia-0 seeds were used as wild type. Mutants used that were previously described are: *hy5-2 hyh* (van Gelderen et al., 2018; Zhang et al., 2017), *hy5-215* (Oyama et al., 1997), *nrt2.1 nrt2.2* (Li et al., 2007) and *chl1-5* (Mounier et al., 2014).

### 2.2 Growth conditions

Plants were grown on either ½MS medium with addition of 1 g/l MES and pH of 5.8 with 0.8 % plant agar (Duchefa), or modified versions of the medium described in Kellermeier et al., 2014 (Table 1), also with the addition of MES and agar. The inserts of the D-root system combined with black paper covers were used to shield the roots from light (Silva-Navas et al., 2015) and the plates were sealed with urgopore tape. The light regime was 16 hours light, 8 hours dark. Photosynthetically Active Radiation (PAR) was 140 μmol/m^2^/s (Philips HPI 400 W), FR light was added using Philips GreenPower LED research modules, far red, 24 Vdc/10 W, 730-nm peak, emitting ~25 μmol/m^2^/s FR light at 20 cm distance. The LEDs were placed at 9 cm height, facing the plates sideways.

**Table 1:** Non-½MS-nutrient media compositions.

|  | normal N<br>(2) mM | low N<br>(0.2) mM | lower N<br>(0.05) mM | NH<sub>4</sub>+NO<sub>3</sub><br>(2) mM | 1.33 NO<sub>3</sub><br>(1.33) mM | +NH<sub>4</sub><br>(2.67) mM | low P<br>(2/0.02) ml |
| --- | --- | --- | --- | --- | --- | --- | --- |
| **macronutrients** |  |  |  |  |  |  |  |
| Potassium nitrate (KNO<sub>3</sub>) | 2.00 | 0.20 | 0.05 | 0.67 | 1.33 | 2.00 | 2.0 |
| Ammonium nitrate (NH<sub>4</sub>NO<sub>3</sub>) |  |  |  | 0.67 |  | 0.67 |  |
| potassium chloride (KCl) |  | 1.80 | 1.95 | 1.33 | 0.67 | 1.95 |  |
| Calcium chloride (CaCl<sub>2</sub> · 2H<sub>2</sub>O) | 0.25 | 0.25 | 0.25 | 0.25 | 0.25 | 0.25 | 0.2 |
| Magnesium chloride (MgCl<sub>2</sub> · 6H<sub>2</sub>O) | 0.25 | 0.25 | 0.25 | 0.25 | 0.25 | 0.25 | 0.2 |
| Magnesium sulphate (MgSO<sub>4</sub> · 7H<sub>2</sub>O) | 0.25 | 0.25 | 0.25 | 0.25 | 0.25 | 0.25 | 0.2 |
| sodium phosphate (NaH<sub>2</sub>PO<sub>4</sub>) | 0.50 | 0.50 | 0.50 | 0.50 | 0.50 | 0.50 | 0.0 |
| sodium chloride (NaCl) | 8.00 | 8.00 | 8.00 | 7.33 | 8.00 | 8.00 | 8.4 |

|  | normal N<br>(2) mM | low N<br>(0.2) mM | lower N<br>(0.05) mM | NH<sub>4</sub>+NH<sub>3</sub><br>(2) mM | 1.33 NO<sub>3</sub><br>(1.33) mM | +NH<sub>4</sub><br>(2.67) mM | low P<br>(2/0.02) ml |
| --- | --- | --- | --- | --- | --- | --- | --- |
| **micronutrients** |  |  |  |  |  |  |  |
| Fe(III)Na-EDTA | 0.04 | 0.04 | 0.04 | 0.04 | 0.04 | 0.04 | 0.0 |
| MnCl<sub>2</sub> · 4H<sub>2</sub>O | 1.80 | 1.80 | 1.80 | 1.80 | 1.80 | 1.80 | 1.8 |
| H<sub>3</sub>BO<sub>3</sub> | 45.00 | 45.00 | 45.00 | 45.00 | 45.00 | 45.00 | 45.0 |
| ZnSO<sub>4</sub> 7H<sub>2</sub>O | 0.38 | 0.38 | 0.38 | 0.38 | 0.38 | 0.38 | 0.3 |
| (NH<sub>4</sub>)<sub>6</sub> Mo<sub>7</sub>O<sub>24</sub> | 0.02 | 0.02 | 0.02 | 0.02 | 0.02 | 0.02 | 0.0 |
| CuSO<sub>4</sub> · 5H<sub>2</sub>O | 0.16 | 0.16 | 0.16 | 0.16 | 0.16 | 0.16 | 0.1 |
| CoCl<sub>2</sub> | 0.01 | 0.01 | 0.01 | 0.01 | 0.01 | 0.01 | 0.0 |
| **Vitamins and organics** |  |  |  |  |  |  |  |
| myo-Inositol 100 mg/l | 0.2775 | 0.2775 | 0.2775 | 0.2775 | 0.2775 | 0.2775 | 0.277 |
| Niacin 0.5 mg/l | 0.0020 | 0.0020 | 0.0020 | 0.0020 | 0.0020 | 0.0020 | 0.002 |
| Pyridoxine · HCl 0.5 mg/l | 0.0012 | 0.0012 | 0.0012 | 0.0012 | 0.0012 | 0.0012 | 0.001 |
| Thiamine · HCl 0.1 mg/l | 0.0001 | 0.0001 | 0.0001 | 0.0001 | 0.0001 | 0.0001 | 0.000 |
| Glycine (recrystallized) 2.0 mg/l | 0.0133 | 0.0133 | 0.0133 | 0.0133 | 0.0133 | 0.0133 | 0.013 |

Temperature was 20 °C and humidity 70 %. Seeds were surface sterilized using chlorine gas (bleach+HCl) for two hours and aerated in a flow cabinet for 15 minutes. Sterilised seeds were sown on one row at 9cm height with 27 seeds on one 12cm square Greiner petri dishes containing agar medium and were the sealed and vernalized at 4 °C for 3-6 days. For growth, plates were placed in white light (WL) first and after one day of germination were placed in either WL or WL+FR. After four days seedlings were transferred from starting plates to new identical plates, but with 5 seedlings per plate. At 8-9 days plates were scanned.

### 2.3 Image acquisition, root phenotyping and data processing

Plates were scanned using an Epson V850 flatbed photonegative scanner at 1200 dpi. Hypocotyl length was analyzed with standard ImageJ tools. Root phenotyping was performed using Smartroot (Lobet et al., 2011). Data was processed with R and statistical analysis was performed with both R and Prism.

### 2.4 Seedling fixation and lateral root primordia analysis

After scanning, seedlings were fixed according to the protocol of Malamy and Benfey, 1997a. Seedlings were mounted in 50% glycerol and slides were sealed with nail polish. Slides were analyzed using a Zeiss Axioskop2 DIC (differential interference contrast) microscope (40X Plan-NEOFLUAR DIC objective) with a Lumenera Infinity1 camera.

### 2.5 RNA extraction and qRT-PCR expression analysis

For gene expression analyses, plants were sown at 16 seeds in a row and kept in the growth conditions mentioned above for 5 days. Between 15 to 19 seedlings were harvested per sample, only root tissues were used for RNA extraction. Four biological replicates were taken per treatment/genotype condition. The Qiagen plant RNeasy kit was used for RNA extraction. First-strand cDNA was made using the Thermo Scientific RevertAid H Minus Reverse Transcriptase, RiboLock RNase inhibitor, and Invitrogen random hexamer primers. RNA input into the cDNA reaction was kept equal within experiments. Primers were designed preferably across introns and for 100-to 150-bp fragments with an annealing temperature of 60°C with primer3plus (http://www.bioinformatics.nl/cgi-bin/primer3plus/primer3plus.cgi). Primers were tested for efficiency using generic Col-0 cDNA at a concentration range of 2.5→40 ng of cDNA per 5 mL reaction. qPCR reagents used were Bio-Rad SYBR-Green Mastermix on 384-well plates in a Life Technologies ViiA7 real-time PCR system. All CT values were normalized against two validated housekeeping genes: *ADENINE PHOSPHORIBOSYL TRANSFERASE1 (APT1) and PROTEIN PHOSPHATASE 2A SUBUNIT A3 (PP2AA3)*. The DDCT method was used to calculate relative expression values (Livak and Schmittgen, 2001). Primer sequences are provided in Table 2.

**Table 2:**
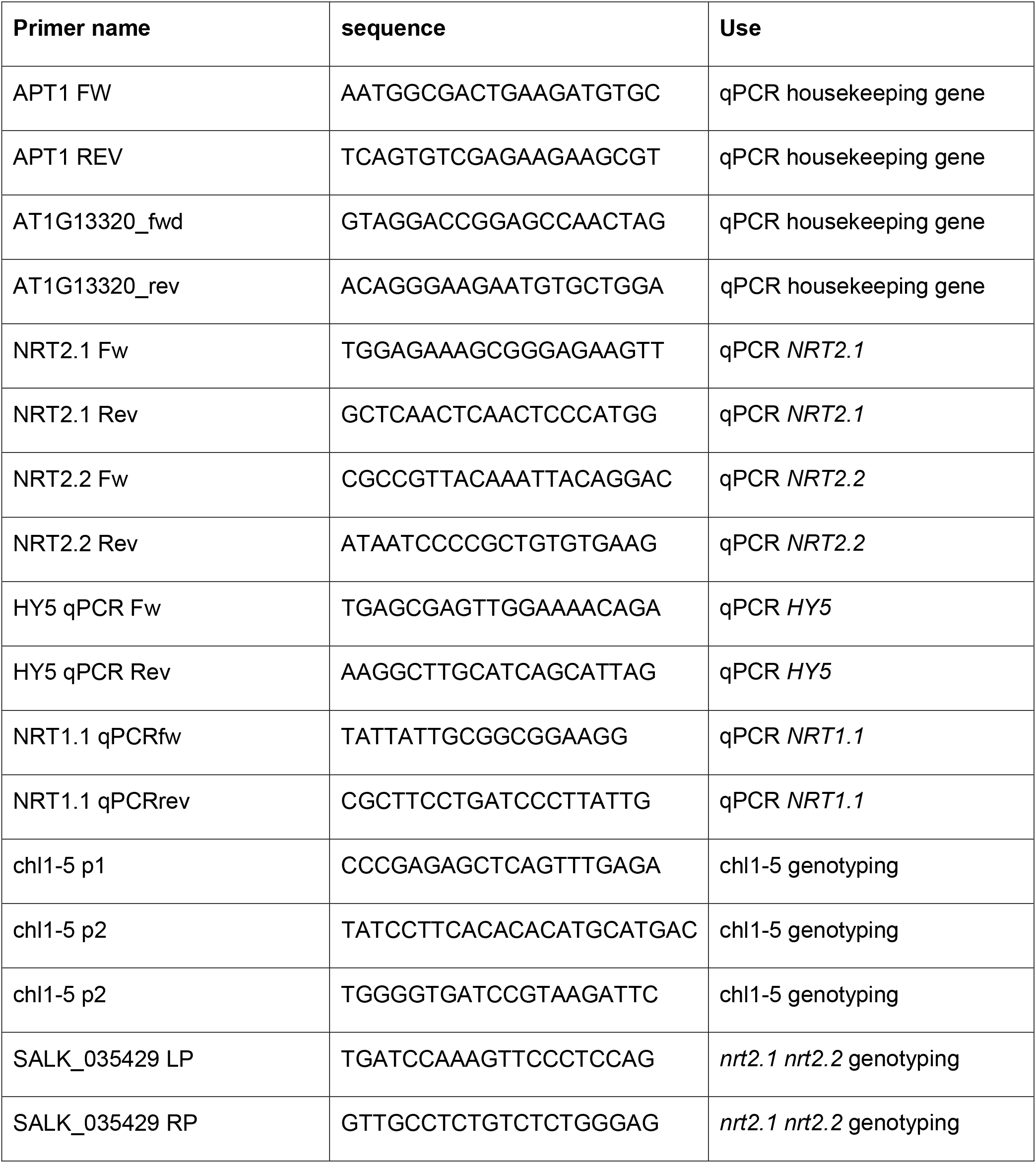
Primers used in study.

## 3 Results

### 3.1 Low Nitrate decreases the response to WL+FR in both the hypocotyl and root

In order to investigate the effect of low nitrate on the root system of *Arabidopsis thaliana*, we used a growth medium with mineral composition as published in (Kellermeier et al., 2014)(Table 1). Ammonium was left out to remove any interfering effects with nitrate signaling (Hachiya and Sakakibara, 2017). Since our previous work (van Gelderen et al., 2018) on the response of the root system to WL+FR was based on plates containing ½ MS we compared the nitrate-only-N medium and ½ MS media with respect to wildtype Col-0 responses to WL+FR. We employed the use of the D-root system (Silva-Navas et al., 2015), in order to grow the root system under physiologically meaningful conditions that avoid light exposure (van Gelderen et al., 2018). Overall, Col-0 wild type responded in a similar manner to WL+FR on the nitrate-only-N medium compared to when these plants were grown ½ MS-containing medium (Figure 1A-D). WL+FR stimulated hypocotyl length (Figure 1A), whereas lateral root density and main root length were reduced (Figure 1B-C). Having confirmed that the nitrate-only-N medium gave similar FR-induced root and shoot architecture phenotypes as on ½ MS we proceeded to investigate the low nitrate response. A ten-fold lower concentration (0.2 mM) compared to control (2 mM) of nitrate resulted in a reduction of hypocotyl elongation in WL+FR when compared to 2 mM nitrate (Figure 1E,H). The reduction of lateral root density and main root length due to WL+FR was lost in the low nitrate condition (Figure 1 F,G,H). These results indicate that a low nitrate medium leads to the loss or reduction of WL+FR-induced changes of root and shoot development. We verified if this is a nitrogen-specific effect, by performing a comparable experiment, but now depleting phosphate. Whereas similarly to low nitrate hypocotyl length was reduced, lateral root density was increased by low phosphate (Figure 1E,F,H), rather than decreased as in low nitrate. Interestingly, WL+FR did not decrease the lateral root density in low phosphate, however it did decrease the main root length (Figure 1F,G). These results show that the specific nutrient status of the medium and/or the plant affects the manner in which root system development integrates with the light spectral composition to which the shoot is exposed.

**Figure 1:**
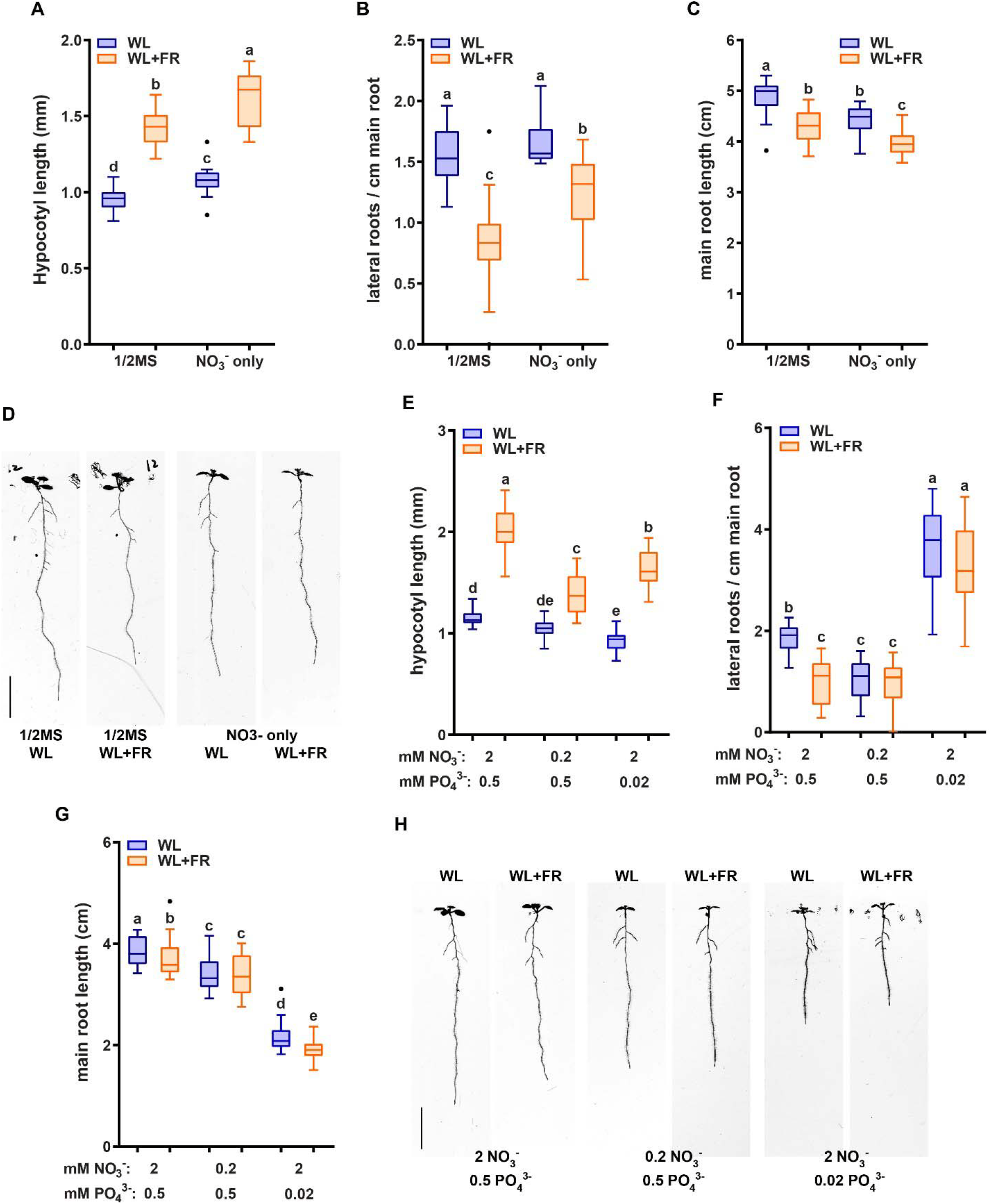
Nutrient shortage decreases the response to low R:FR in both the hypocotyl and root. (A-D) Seedlings (Col-0) were grown in D-root plates (Silva-Navas et al., 2015), with WL and WL+FR light conditions described in (van Gelderen et al., 2018) and supplied with either regular 1/2MS or with the control medium of (Kellermeier et al., 2014), which does not contain ammonium as a nitrogen source. (A) Hypocotyl length, (B) Main root length, (C) Lateral root density. (D) Representative seedlings for data in A,B and C. (E-H) Analysis of 8-day-old seedlings grown in either control (2mM NO3^−^), low nitrate medium (0.2mM), or low phosphate (0.2 mM). (E) Hypocotyl length, (F) Main root length, (G) Lateral root density. (H) Representative seedlings for data in A,B and C. Letters denote statistically significant groups based on a mixed model 2-way ANOVA with posthoc tukey test (p<0.05). Scale bar = 1cm.

### 3.2 NRT2.1/2.2 and HY5 HYH are required for the combined WL+FR and nitrate response

Previous work identified that root development of the *hy5 hyh* double mutant is unresponsive to WL+FR (van Gelderen et al., 2018). Furthermore, HY5 regulates nitrate uptake via the transcriptional control of the nitrate transporter gene *NRT2.1* (Chen et al., 2016; Jonassen et al., 2009, 2008). NRT2.1 is part of the high affinity nitrate uptake pathway and NRT2.1 inhibits the initiation of lateral root primordia during nitrate starvation (Remans et al., 2006; Little et al., 2005). Therefore, we tested both *hy5 hyh* and *nrt2.1 nrt2.2* mutants in WL or WL+FR conditions on normal and low nitrate media (a single T-DNA knocks out both *nrt2.1* and *2.2* (Li et al., 2007)). WL+FR again led to a decrease in main root length in Col-0, which was abolished in low nitrate (Figure 2A,C). Both *nrt2.1 nrt2.2* and *hy5 hyh* lacked the WL+FR-induced main root length decrease, but did show a strong main root length reduction upon exposure to low nitrate (Figure 2A,D,E). Col-0 lateral root density on normal nitrate was reduced by WL+FR, however this did not occur in *nrt2.1 nrt2.2* (Figure 2B), while low nitrate strongly reduced *nrt2.1 nrt2.2* lateral root density (Figure 2B). Even more striking was the fact that the lateral root density of *hy5 hyh* did not significantly change in any of the conditions (Figure 2B). The hypocotyl elongation responses to WL+FR of both *nrt2.1 nrt2.2* and *hy5 hyh* mutants were similar in trend to Col-0, however *nrt2.1 nrt2.2* was slightly more sensitive to low nitrate and *hy5 hyh* has a much longer hypocotyl length to start out. (Figure S1A). To test the limits of nitrate depletion further we grew the same mutants on a lower concentration of nitrate (0.05 mM). Both Col-0 and *nrt2.1 nrt2.2* lateral root density were severely reduced by this depletion, however the lateral root density of *hy5 hyh* was only slightly affected (Figure S1B). These results confirm the enhanced sensitivity of *nrt2.1 nrt2.2* to low nitrate conditions, but also indicate a reduced sensitivity of the *hy5 hyh* mutant to low nitrate. Furthermore, both mutants lacked the lateral root density response to WL+FR, showing that it is likely they are both involved in mediating the response to WL+FR.

**Figure 2:**
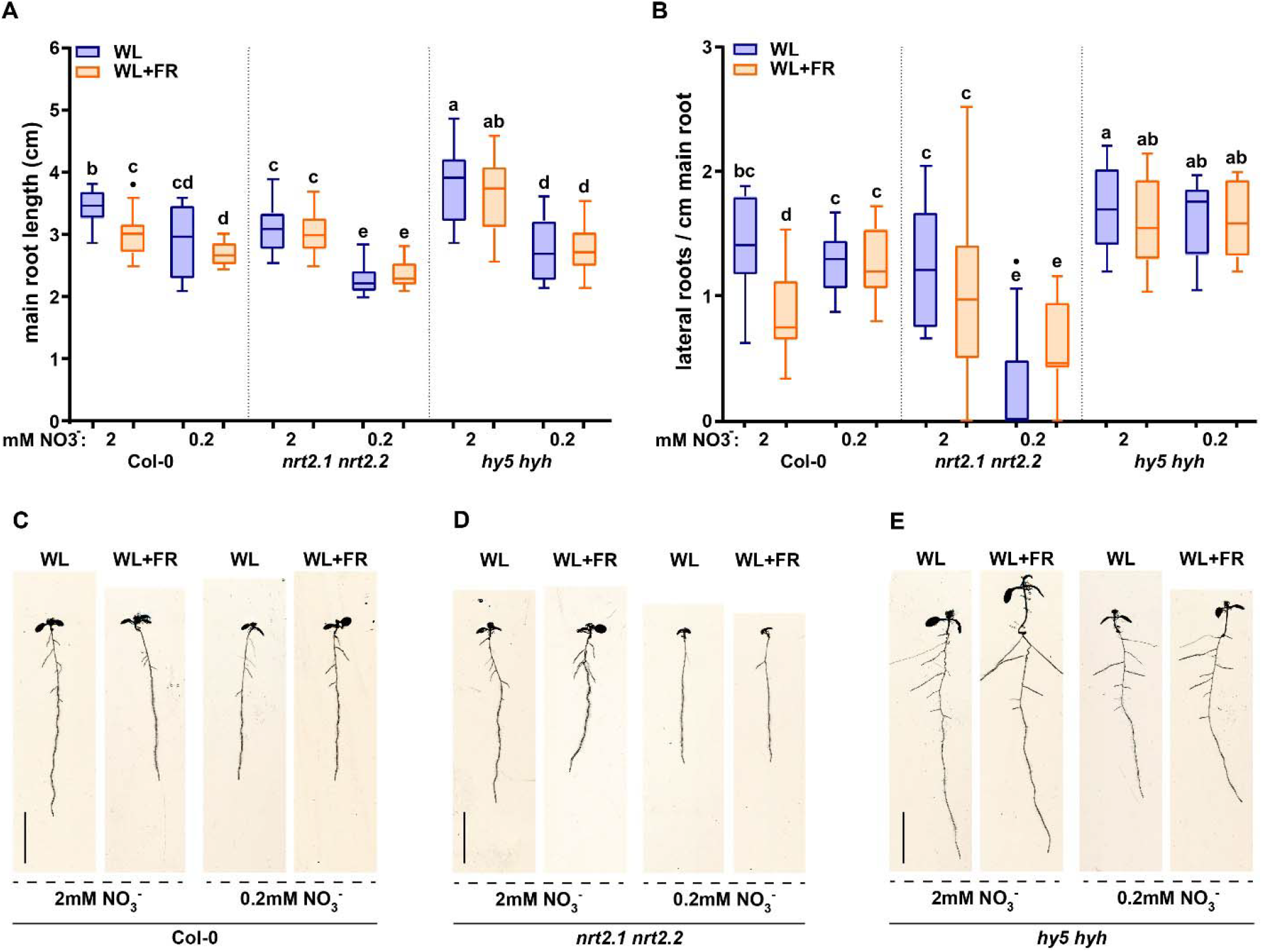
*NRT2.1/2.2* and *HY5 HYH* are required for the combined low R:FR & nitrate response. Wild type Col-0, *hyh5 hyh* and *nrt2.1 nrt2.2 s*eedlings grown for eight days in either WL or WL+FR on normal or low nitrate media. (A) Main root length, (B) Lateral root density. (C,D,E) Representative seedlings for data in A and B. Letters denote statistically significant groups based on a mixed model 2-way ANOVA with posthoc tukey test (p<0.05). Scale bar = 1cm.

### 3.3 Low nitrate medium affects lateral root primordia development

The results presented here and from various other works show that lateral root growth is affected by low nitrate conditions (Kellermeier et al., 2014; Gruber et al., 2013; Bouguyon et al., 2016). It is also known that in certain conditions *NRT2.1* can inhibit lateral root initiation (Little et al., 2005; Remans et al., 2006). To get more insight into what developmentally occurs with lateral root primordia (LRP) growth - whether there is a decrease in initiation, emergence, or a mid-development arrest – we analyzed the seedlings shown in Figure 2 for the frequency of lateral root primordia stages, according to the classification of Malamy and Benfey, 1997 (Figure 3). Col-0 seedlings exposed to WL+FR and grown on 2 mM nitrate had an increase in stage 1+2 and stage 5+6 lateral root primordia when compared to WL-grown seedlings, while the emerged primordia (7+E) were decreased (Figure 3A). This result was similar to a previously published experiment performed on ½ MS-containing plates (van Gelderen et al., 2018) and indicates that primordia are formed, but do not fully develop into lateral roots. Col-0 seedlings on low nitrate (0.2 mM) plates had no LRP stage frequency differences between WL and WL+FR (Figure 3A). Strikingly, *nrt2.1 nrt2.2* mutant seedlings did not have any significant differences between treatments, indicating that both the effect of low nitrate and WL+FR on LRP development are dependent upon *nrt2.1 nrt2.2* (Figure 3B). The *hy5 hyh* LRP stages did have significant differences, which overall appeared to be opposite to those in Col-0. In the *hy5 hyh* genotype, WL+FR led to more 7+E stages in normal and low nitrate, while LRP stage 1+2 and 3+4 frequency was reduced (Figure 3C). However, the effect of low nitrate on *hy5 hyh* LRP stages was relatively minor, when compared to Col-0. It is surprising that there were significant changes in late lateral root primordia stages in *hy5 hyh*, since the lateral root density hardly changed between the four conditions (Figure 2C). In *hy5 hyh* there was not a significant difference in the total number of primordia between WL and WL+FR (Figure S2), thus it is possible that these extra primordia have not yet resulted in a visually changed lateral root outgrowth. Overall, these results show that low nitrate removes any WL+FR effect on lateral root primordia in Col-0 and that *nrt2.1 nrt2.2* is unresponsive to both low nitrate and WL+FR, while *hy5 hyh* has a distinctly different response than Col-0.

**Figure 3:**
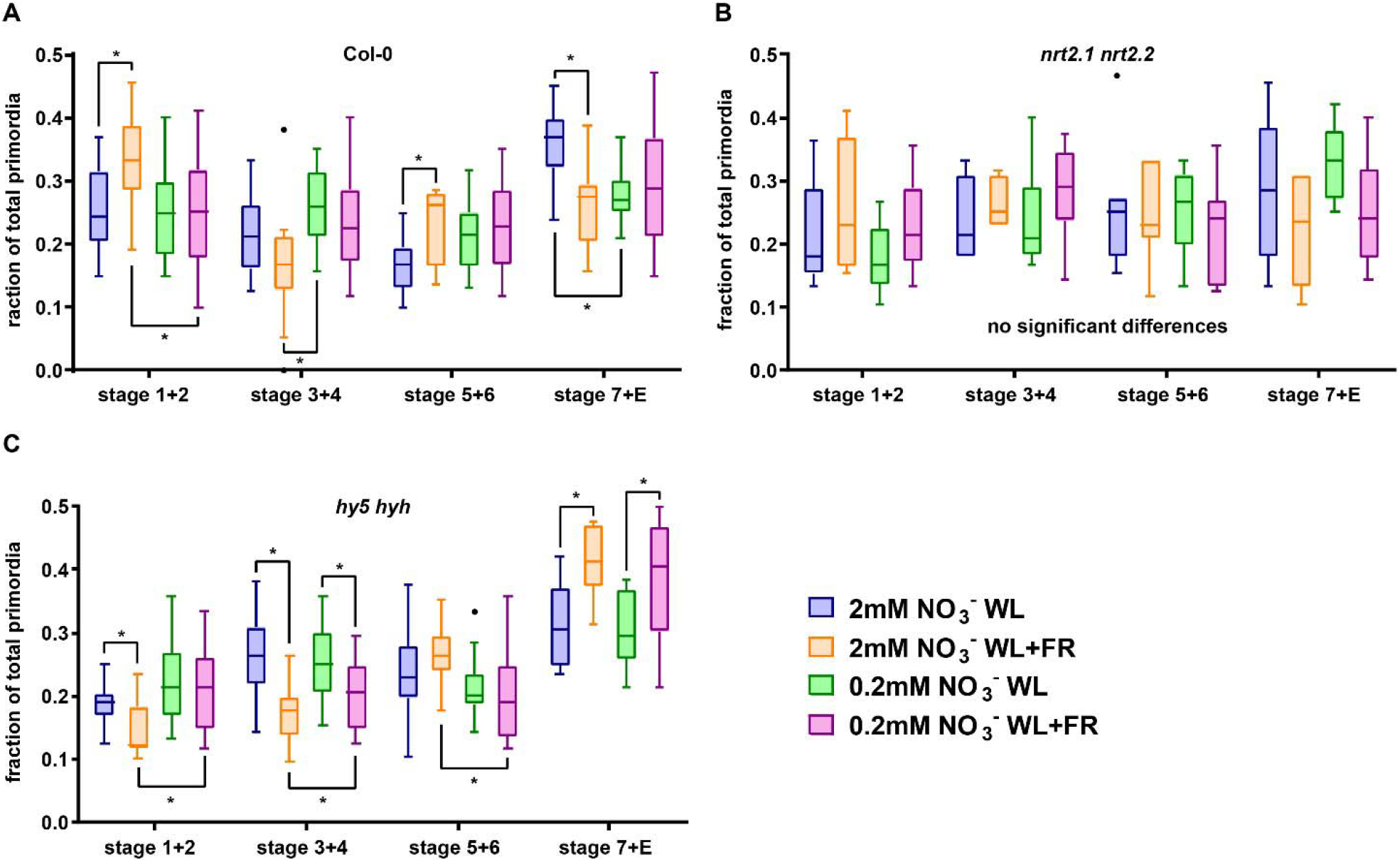
Low nitrate conditions regulate lateral root primordia development. Lateral root primordia (LRP) data from 8-day old seedlings from experiment in Figure 2. Stages are counted according to(Malamy and Benfey, 1997) and grouped in pairs of two going from early (1+2) to late (7+Emerged) development. (A) Col-0, (B) *nrt2.1 2.2*, (C) *hy5 hyh. Statistics*: mixed model 2-way ANOVA with post hoc Tukey test within the stage groups (*p<0.05).

### 3.4 *HY5* and WL+FR regulate *NRT2.1* expression

HY5 is able to promote transcription of *NRT2.1* and it is also upregulated by low nitrate levels (Chen et al., 2016; Okamoto et al., 2003). Since HY5 is crucial for the root response to shoot-perceived WL+FR (Figure 2), we were interested in investigating if WL+FR can also affect *NRT2.1* expression. We performed a time-course qRT-PCR experiment seedling root material during the 16 hours photoperiod to map when the response of *HY5* and *NRT2.1* to WL+FR was strongest (Figure 4A,B). We observed that during the day *NRT2.1* expression kept rising and that, except at 12 hours post-dawn, WL+FR led to an additional increase in *NRT2.1* expression (Figure 4A). *HY5* expression peaked at 4 and 8 hours into the day, at which time it was also upregulated by WL+FR (Figure 4B). Next, we grew Col-0 and *hy5 hyh* seedlings on normal or low nitrate plates, and in WL or WL+FR. *NRT2.1* was upregulated in Col-0 WL+FR treated seedlings (Figure 4C). We did not observe the expected strong increase of *NRT2.1* transcription due to the low nitrate levels, however that can be explained by the duration of the low nitrate treatment (5 days), since upregulation of *NRT2.1* by low nitrate is a transient effect and decreases slowly after 1 day (Okamoto et al., 2003). Importantly, in the *hy5 hyh* mutant background *NRT2.1* transcription was upregulated and the increase in WL+FR was absent, however the increase due to low nitrate was still present (Figure 4C). The observed elevated expression of *NRT2.1* in the *hy5 hyh* mutant is in contrast with the previously shown stimulation of this gene by HY5 (Chen et al., 2016). We performed a similar experiment with another *hy5* mutant, *hy5-215*, on normal nitrate medium, and the result was the same (Figure S3A). However, we observed that when we grew *hy5-215* on ½ MS medium, a strong decrease of *NRT2.1* expression was observed (Figure S3B). Therefore, we hypothesized that the addition of ammonium might be a crucial element in regulating *NRT2.1* expression through HY5. To test this, we made a medium with 2 mM N consisting of 1.33 mM NO_3_^−^ and 0.67 mM NH_4_^+^, a nitrate/ammonium ratio that is very similar to ½ MS. When we repeated the experiment with the addition of this combined nitrate-ammonium we observed a strong decrease in *NRT2.1* expression on the combined nitrate-ammonium medium, both in wild type, and *hy5 hyh* (Figure 4D). Again, we observed an increase in *NRT2.1* expression due to WL+FR and in the *hy5 hyh* mutant background (Figure 4D). In this experiment there was no increase in the WL+FR & low nitrate condition. In addition, we tested the expression of the close homologue *NRT2.2*, which displayed a strong response to WL+FR and a very strong response to low nitrate (Figure 4E). Its expression was higher in the *hy5 hyh* background, however in this mutant *NRT2.2* did still respond to WL+FR and low nitrate, indicating that *NRT2.2* is regulated in a somewhat different manner than *NRT2.1*. *NRT2.2* expression was decreased in the combined nitrate-ammonium medium to almost undetectable levels (Figure 4E). We tested the expression of *HY5* on the same material and observed an increase in expression due to WL+FR and interestingly, also an increase due to low nitrate (Figure 4F). In both low nitrate and combined nitrate-ammonium medium the expression of *HY5* did not change due to WL+FR (Figure 4F), while *HY5* expression was not changed in the *nrt2.1 nrt2.2* mutant (Figure 4G). These results show that shoot-perceived FR light and low nitrate leads to increased expression of *NRT2.1, NRT2.2* and *HY5*. However, in low nitrate the upregulation of these genes due to shoot-perceived FR is less. In the *hy5 hyh* mutant, *NRT2.1* and *NRT2.2* are upregulated, indicating a negative effect of *HY5* on their transcription, however this effect can be fully masked by the addition of ammonium. Importantly, in the *nrt2.1 nrt2.2* mutant, *HY5* expression is not changed, indicating that NRT2.1 likely acts downstream, and not upstream, of HY5.

**Fig. 4.**
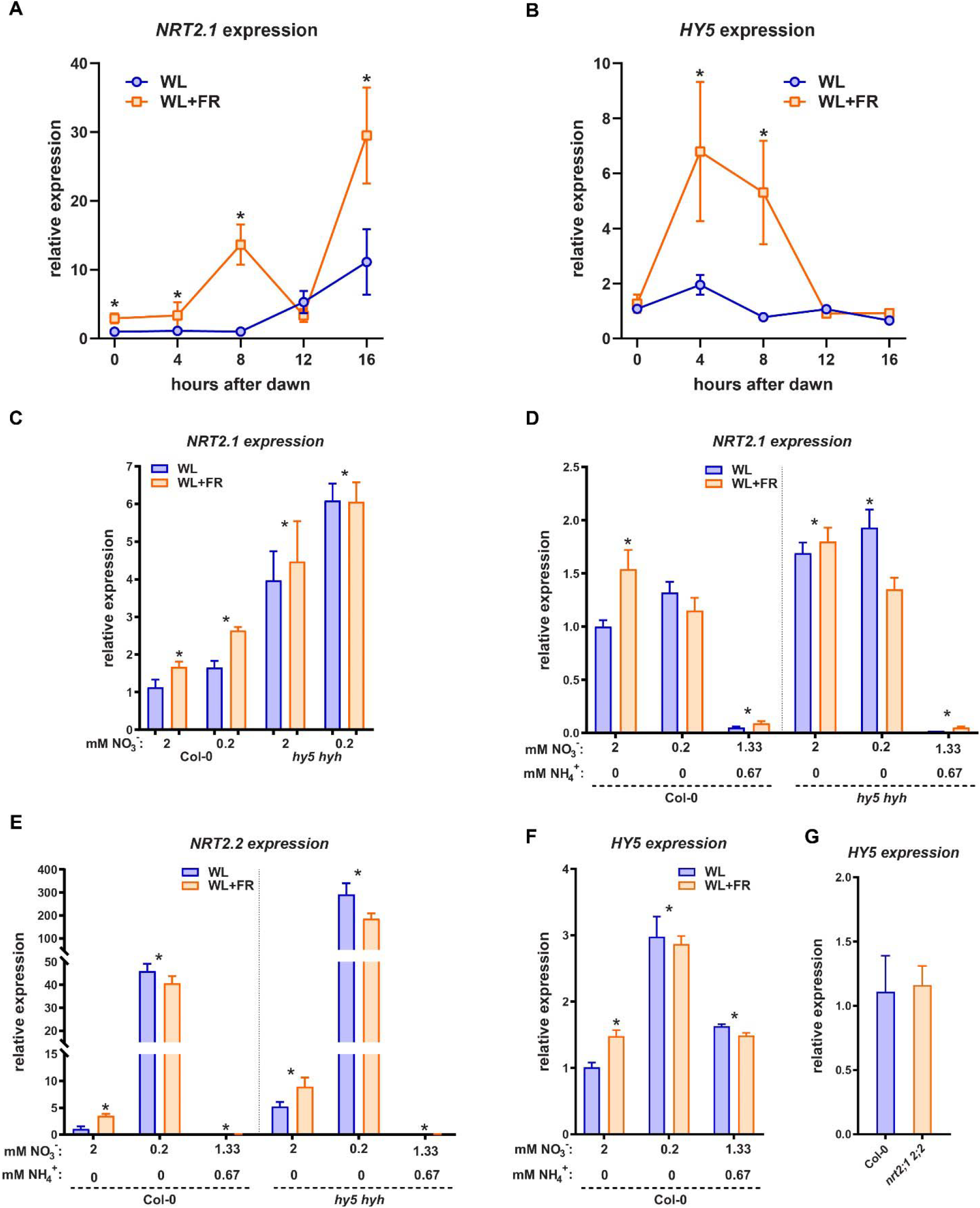
*HY5* and WL+FR regulate *NRT2.1* expression. (A,B) Time-course qPCR experiment using 5-day old seedling root material harvested between 0 – 16 hours post-dawn, grown either in WL or WL+FR. WL+FR increases *NRT2.1 2.2* expression in all timepoints bar 12hrs PD. *HY5* expression is increased by WL+FR at 4 and 8 hours PD. (C) qPCR expression analysis of *NRT2.1* using RNA of 5-day old seedling roots, with combined WL/WL+FR and low N / normal N treatments. (D,E,F) qPCR expression analysis similar to (C), but with the addition of a mixed nitrate/ammonium medium (NO ^−^ NH ^+^), (D): *NRT2.1* (E): *NRT2.2* (F): *HY5*. *HY5* expression was not detectable in the *hy5 hyh* mutant. (G) qPCR expression analysis similar to (C), of *HY5* on normal (2 mM) nitrate medium in Col-0 and *nrt2.1 2.2*. (A,B) *p<0.05 WL vs WL+FR with a two-way ANOVA plus post-hoc tukey test, (C-F) *p<0.05 vs 2mM Col-0 WL with a one-way ANOVA plus post-hoc tukey test.

### 3.5 Ammonium addition masks WL+FR effect on root development

Addition of ammonium as a nitrogen source led to a very strong decrease in *NRT2.1* and *NRT2.2* expression. This prompted us to investigate if the addition of ammonium had significant effects on the root developmental response to WL+FR. Ammonium can stimulate lateral root initiation and directly promote lateral root emergence, while ammonium-dependent signaling can decrease part of the primary low nitrate response (Hachiya and Sakakibara, 2017; Lima et al., 2010; Meier et al., 2020). The addition of ammonium had little effect on hypocotyl elongation (Figure S4A). Interestingly, the addition of ammonium led to the loss of difference in lateral root density between WL and WL+FR as seen in Col-0 on nitrate-only-N medium (Figure 5A,C). On nitrate-only-N medium, the *nrt2.1 nrt2.2* mutant had a reduced lateral root density compared to Col-0 WL, but the addition of ammonium removed this effect and led to increased lateral root density (Figure 5A). On the combined ammonium-nitrate medium there was no difference in main root length between WL and WL+FR (Figure 5B). To ensure that this effect was due to the ammonium addition and not to the concomitant reduction of available nitrate, we included an additional control where we added 0.67 mM ammonium in addition to the 2 mM nitrate and a medium with 1.33 mM nitrate as only N-source (Figure S4B, Table 1). Again, we saw that low (0.2 mM) nitrate led to a lower lateral density without a WL+FR-induced reduction and that the combined nitrate-ammonium medium had a higher lateral root density that was not affected by WL+FR. Adding 0.67 mM ammonium to 2 mM or to 1.33 mM nitrate gave the same results, indicating that in the combined-N media, it is the variation in ammonium, not nitrate, that affects the phenotypes (Figure S4B). The mild depletion of nitrate (1.33 mM) was comparable to 2 mM nitrate; the lateral root density in WL was similar and WL+FR led to a reduction in lateral root density (Figure S4B).

**Figure 5:**
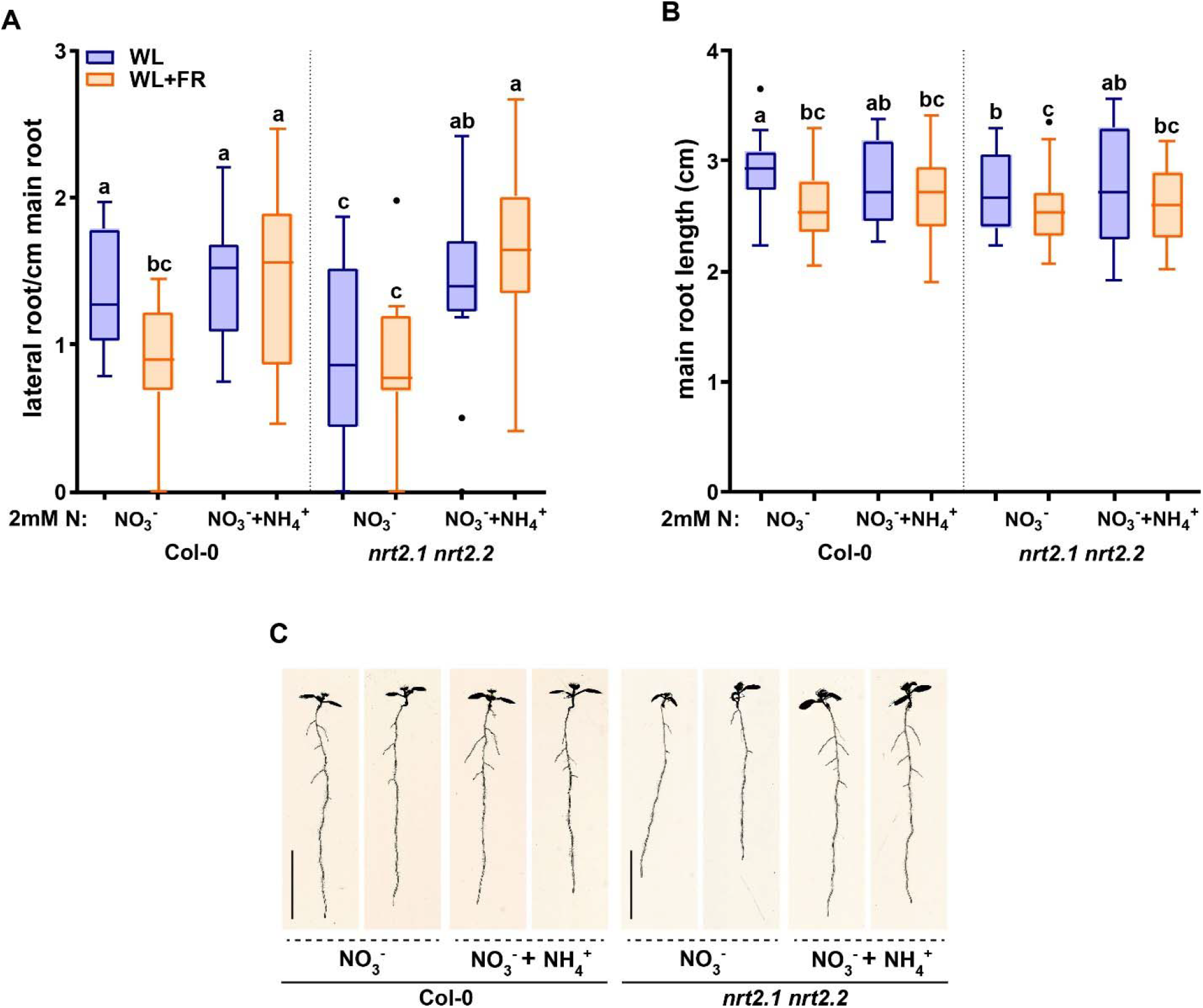
Replacing 1/3 of nitrate with ammonium can remove effect of WL+FR on root development and bypass the *nrt2.1 2.2* root phenotype. Seedlings of Col-0 and *nrt2.1 nrt2.*2 were grown on combined nitrate-ammonium consisting of 1.33 mM NO_3_ ^−^, and 0.67 NH_4_ ^+^. The rest of the experiment was performed according to Figures 1 & 2. (A) hypocotyl length, (B) main root length, (C) lateral root density and (D) representative 8d. old seedlings. Letters denote statistically significant groups based on a mixed model 2-way ANOVA with posthoc tukey test (p<0.05). Scale bar = 1cm.

On the combined nitrate-ammonium medium we observed a decrease in *NRT2.1* expression (Figure 4D). It is known that *NRT2.1* expression in ammonium-containing medium is derepressed in the *NRT1.1* mutant *chl1-5* (Muños et al., 2004; Bouguyon et al., 2015). Therefore, we tested the *chl1-5* mutant on normal and low nitrate. *chl1-5* had a similar lateral root density as Col-0 in WL normal nitrate, however, in WL+FR it had a higher lateral root density, and not a lower lateral root density (Col-0) or no difference between WL and WL+FR (*nrt2.1 nrt2.2*) (Fig. S4B). This suggests that the response of the *nrt2.1 nrt2.2* mutant on nitrate only-N media is specific to these transporters and not generic to any nitrate transporter. Interestingly the expression of *NRT1.1* could also be induced by WL+FR, and future studies would be required to investigate how this would functionally integrate with the WL+FR response. Our findings on nitrate versus combined nitrate-ammonium media show the importance of the nitrogen source, as compared to the amount of available nitrogen. These findings also indicate that the WL and WL+FR effects we observe in normal and low nitrate and the differences due to ammonium are likely due to changes in signaling rather than a limiting effect of the available nitrogen for growth. Lateral root density of *nrt2.1 nrt2.2* on the combined nitrate-ammonium medium was similar between WL and WL+FR and similar to Col-0 on the same medium, indicating that the ammonium-related root architectures are independent of *nrt2.1 nrt2.2* and insensitive to supplemental FR.

## 4 Conclusion and Discussion

Low R:FR signaling indicates nearby vegetation and induces complex developmental outputs in shade intolerant plants. In the shoot of young seedlings the relative increase in FR light leads to increased elongation, while in the root it leads to a reduction in root elongation and lateral root formation (van Gelderen et al., 2018). Here we have shown that a reduction in nitrate levels reduces the WL+FR response of the shoot and the root. However, it is only the shoot that detects the FR light in the experiments presented here, because we made use of the D-root system. HY5 appears to acts as a shoot-to-root signal that links the perception of FR light in the shoot to the root possibly via shoot-to-root transport, or via unknown intermediates (van Gelderen et al., 2018; Chen et al., 2016). Both *nrt2.1 nrt2.2* and *hy5 hyh* mutants lack a lateral root density response to WL+FR. However, these mutants did respond differently to WL+FR and also to low nitrate. It was striking that the *hy5 hyh* mutant root development was insensitive to very low nitrate levels, irrespective of the R:FR light ratio. This indicates that HY5 can play a central role in the adaptation of the root system to low nitrate, as well as to shoot-perceived WL+FR. The fact that on normal nitrate *nrt2.1 nrt2.2* did not show a reduction in lateral root density upon supplemental FR exposure shows that it is involved in this response when nitrate conditions are not limiting. Overall, we can conclude that it is likely that *NRT2.1* has a positive effect on lateral root development, since the *nrt2.1 2.2* mutant had a slightly lower lateral root density compared to Col-0 and that HY5 has a negative effect, since the mutant had a higher lateral root density. It is not surprising that the *nrt2.1 nrt2.2* mutant is more sensitive to low nitrate, since it probably has an impaired nitrate uptake (Remans et al., 2006). Therefore, it is a distinct possibility is that the low nitrate insensitivity of *hy5 hyh* (Figure S1B) is due to the increase in *NRT2.1* transcript, thereby tentatively enhancing nitrate uptake, making it less sensitive to nitrate reduction in the medium. However, we cannot conclude this for the whole seedling, since low R:FR-induced hypocotyl elongation in *hy5 hyh* is still affected by low nitrate.

It is very interesting that low nitrate led to an increase in *HY5* expression. *HY5* overexpressing lines have a reduced lateral root density and do not have a reduced lateral root growth due to shoot perceived WL+FR (Sibout et al., 2006; van Gelderen et al., 2018). Our qPCR data are consistent with the suggestion that NRT2.1 acts downstream of HY5 in the lateral root density WL+FR response. Furthermore, *NRT2.1* was upregulated in the root when the shoot was in WL+FR and this upregulation was dependent upon *HY5*. This is in accordance with our finding that *NRT2.1* is important for the response to WL+FR. Thereby we suggest a model (Figure 6) where low R:FR induces and stabilizes HY5 in the shoot, after which it could be transported to the root, where through a positive feedback mechanism it stimulates its own transcription (Zhang et al., 2017). This leads to repression of lateral root development, directly via for example *ARF19* (van Gelderen et al., 2018), and indirectly via repression of *NRT2.1*. In low nitrate conditions, *NRT2.1* is upregulated at first, but this effect is transient (Okamoto et al., 2003), which explains the relatively mild induction of *NRT2.1* observed in plants exposed to low nitrate for 5 d (Figure 4C). However, low nitrate also stimulates *HY5* expression (Fig. 4F), which would likely reduce lateral root density (Sibout et al., 2006; van Gelderen et al., 2018).

**Figure 6:**
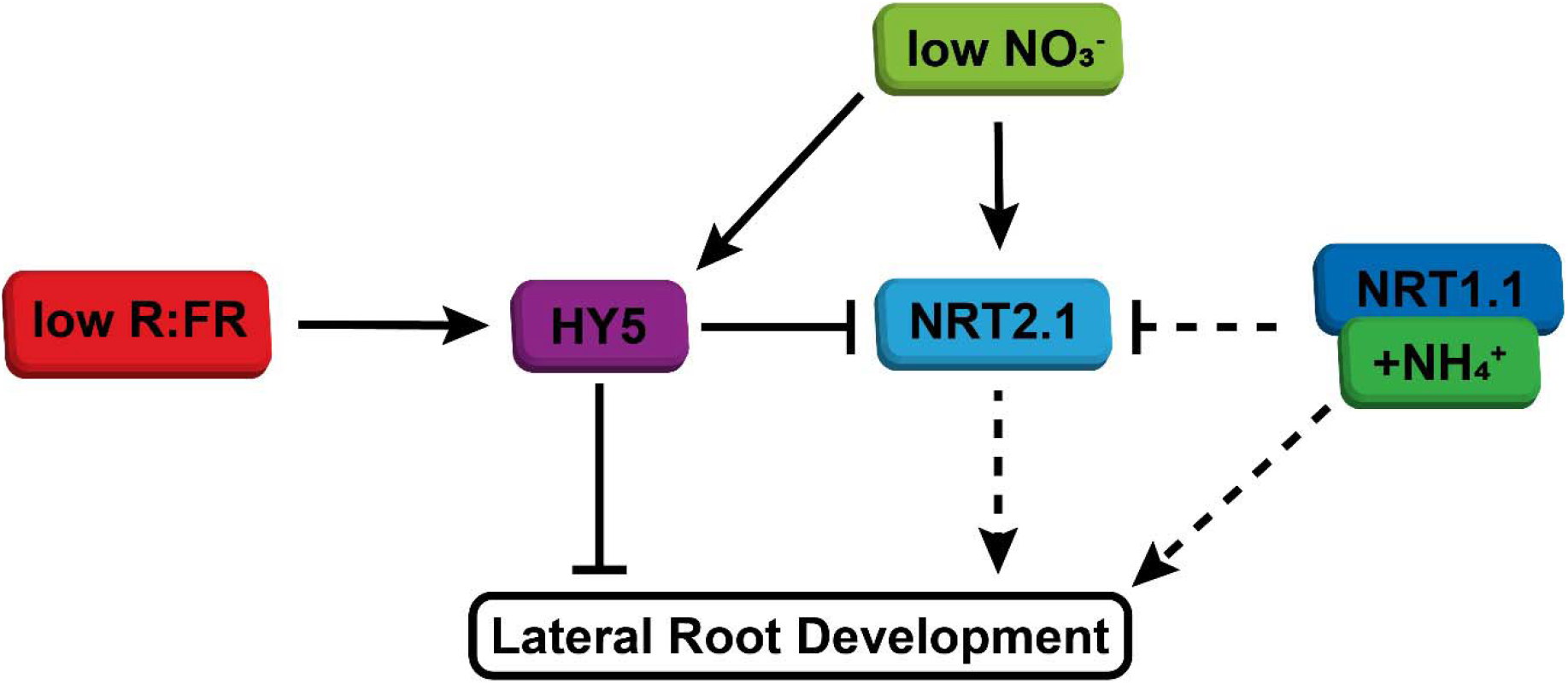
Simplified model of integration of shoot-perceived low R:FR and low nitrate availability on lateral root development. WL+FR enhances HY5 expression, stabilization and transport. In normal nitrate conditions this leads to reduced lateral root development, both due to direct effect of HY5 on emergence, but also due to repression of NRT2.1, which has a positive effect on lateral root development. When nitrate availability is low, both *HY5* expression and *NRT2.1* expression is enhanced, which results in further repression of lateral root development through HY5. Ammonium, possibly via NRT1.1, bypasses low R:FR signaling through its direct effect on lateral root development and also via its repression of NRT2.1 induction.

From the mutant analysis we conclude that *NRT2.1* has a positive effect on root growth. The nutrient context is crucial for the effect of the *nrt2.1* mutation (Little et al., 2005; Remans et al., 2006). We found this as well, since the positive effect of *NRT2.1* on lateral root formation was only true for nitrate-only-N media, as the *nrt2.1 nrt2.2* mutant had a wild type lateral root density and main root length when ammonium was used in addition to nitrate. The simplest explanation for this result is that ammonium directly stimulates lateral root outgrowth via acidification of the apoplast, leading to increased pH-dependent auxin transport (Meier et al., 2020). This could also mask the strong inhibitory effect of ammonium on *NRT2.1* transcription (Muños et al., 2004; Bouguyon et al., 2015), since negative effects of reduced NRT2.1 on lateral root development would be counteracted by pH-driven auxin transport. Increased *NRT2.1* expression in the *hy5 hyh* mutant is opposite of the result obtained by Chen et al., 2016, but this is only true in nitrate-only-N media. When tested on medium with supplemented ammonium, or ½ MS medium, our data were consistent with Chen et al., 2016, who also used MS medium. These expression results with combined nitrate-ammonium media do highlight that the effect of HY5 on the transcription of downstream genes is not always black and white and also relies upon other factors (Burko et al., 2020).

In this study we tried to answer the question how a plant can integrate different signals coming from the shoot and the root. When a plant is competing for available light it is important to adjust its development. However, it is possible that it will only do so when it can afford to. In other words, only when there are enough nutrients will the plant sacrifice some development of the root system. These carbohydrates are very useful for investing in short-term shoot growth. However, we show here that under nutrient-depleted conditions, root system development does not respond to shoot-detected FR anymore, probably safe-guarding nutrient uptake possibilities.

Concluding, we have shown that nitrate levels can modulate the response to low R:FR-induced stimuli of neighbour competition and that this integration involves the HY5 transcription factor and that the NRT2.1 nitrate transporter play an important role in this integration. It is not yet clear exactly how NRT2.1 affects lateral root development. It could be that it is due to its ability as a nitrate transporter, but possibly also as an active signaling component. Since NRT2.1 acts as a transporter/receptor, it has also been put forward that NRT2.1 could affect lateral root outgrowth via differential expression of aquaporins (Li et al., 2016). In this way, NRT2.1 could locally stimulate water uptake and turgor pressure of the cells around the lateral root primordium, affecting lateral root emergence. Elucidating the exact mechanisms through which NRT2.1 regulates lateral root development responses to nitrate and light are important questions for future research.

## 6 Conflict of Interest

The authors declare that the research was conducted in the absence of any commercial or financial relationships that could be construed as a potential conflict of interest.

## 7 Author Contributions

K.v.G. and R.P. designed the experiments and wrote the manuscript. K.v.G., C.K. and P.L. performed and analysed experiments.

## 8 Funding

This research was funded by the Netherlands Organisation for Scientific Research, open competition grant 823.01.013 and Vici grant 865.17.002 to R.P., a scholarship of government sponsorship for overseas study, Taiwan, admission number 0991167-2-UK-004 to C.K.

## 9 Acknowledgments

The authors would like to thank Dr. Sandrine Ruffel for providing the *chl1-5* mutant and for suggestions on the direction of the project. Special thanks go out to Jannetje Kooij and Koen Bensink for practical help during the project.

---

**Figure S1.**
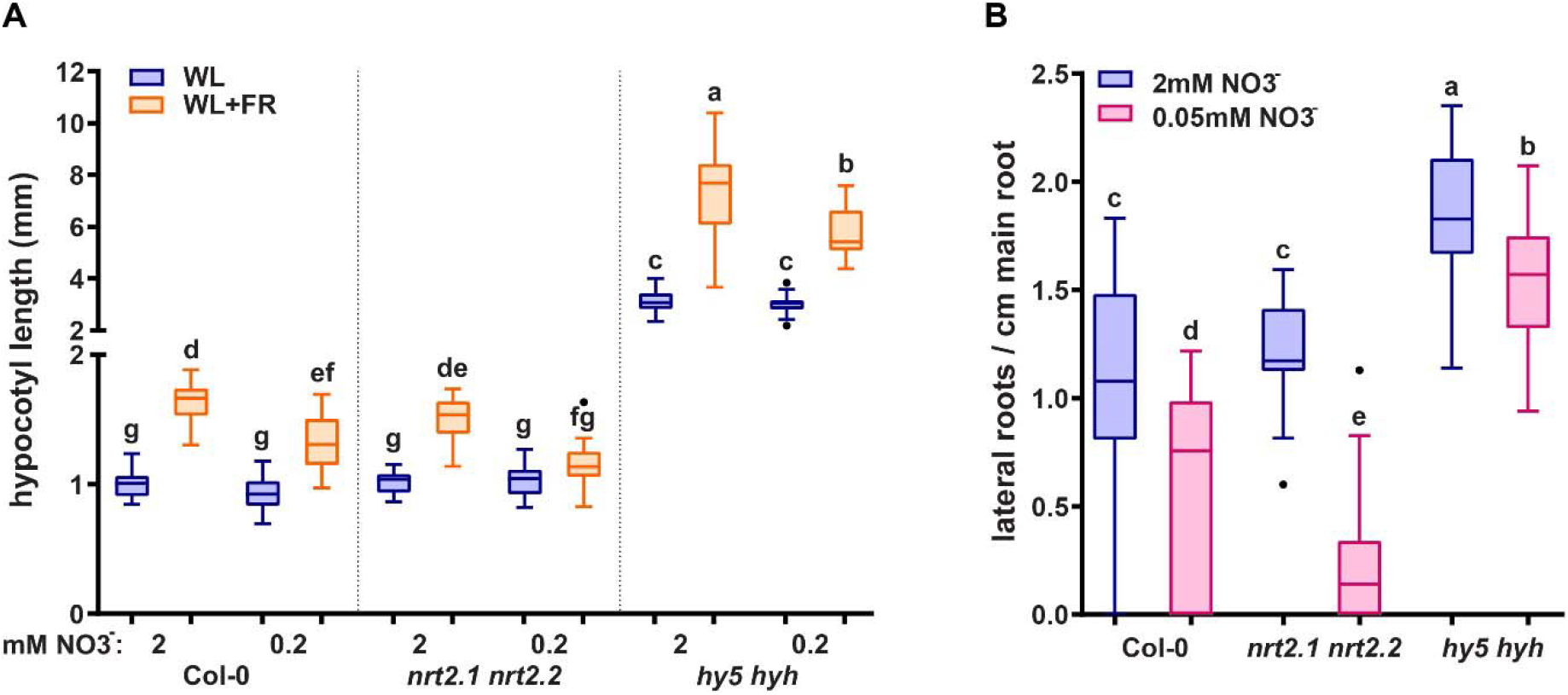
(A) Hypocotyl length of eight day old seedlings grown in WL on normal (2 mM) or low (0.2 mM) nitrate. (B) Lateral root density of eight day old seedlings grown in WL on normal (2 mM) or very low (0.05 mM) nitrate. Letters denote statistically significant groups based on a mixed model 2-way ANOVA with posthoc tukey test (p<0.05). Scale bar = 1cm.

**Figure S2.**
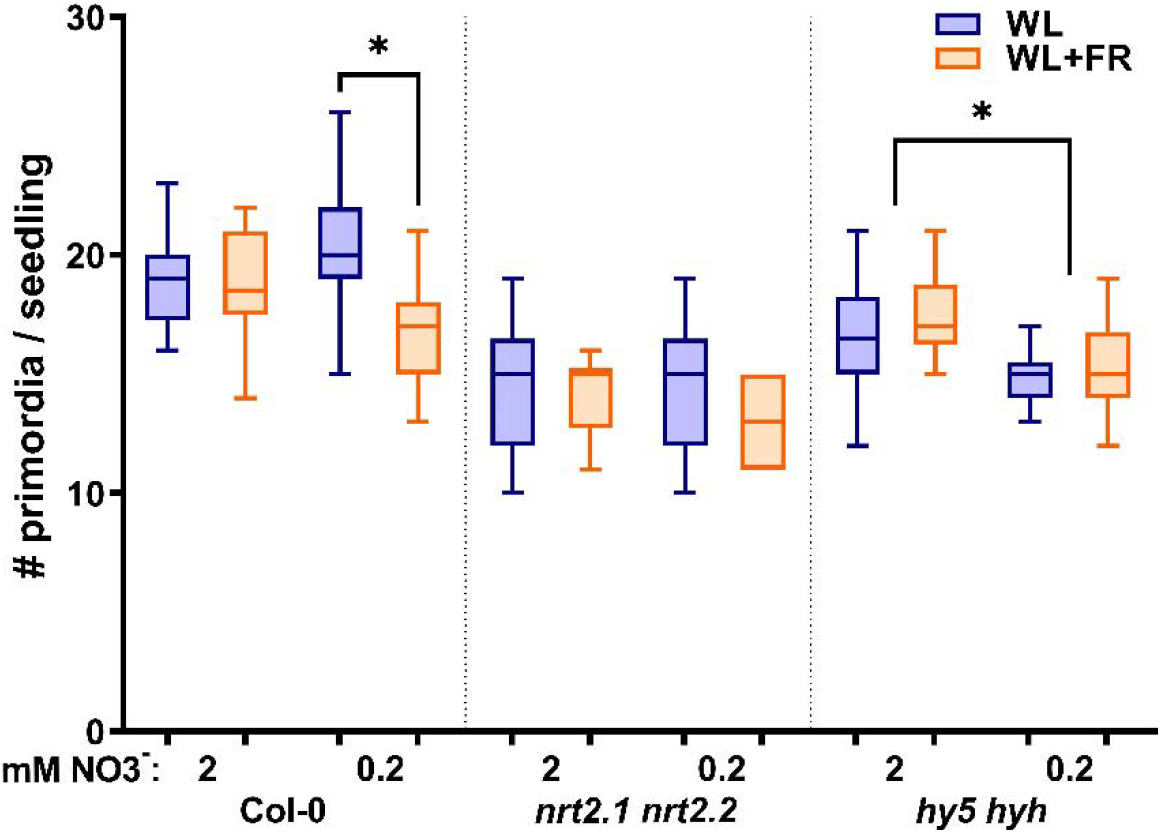
Total number of primordia per seedling from the experiment shown in figure 3. *Statistics*: mixed model 2-way ANOVA with post hoc Tukey test within the stage groups (*p<0.05).

**Figure S3.**
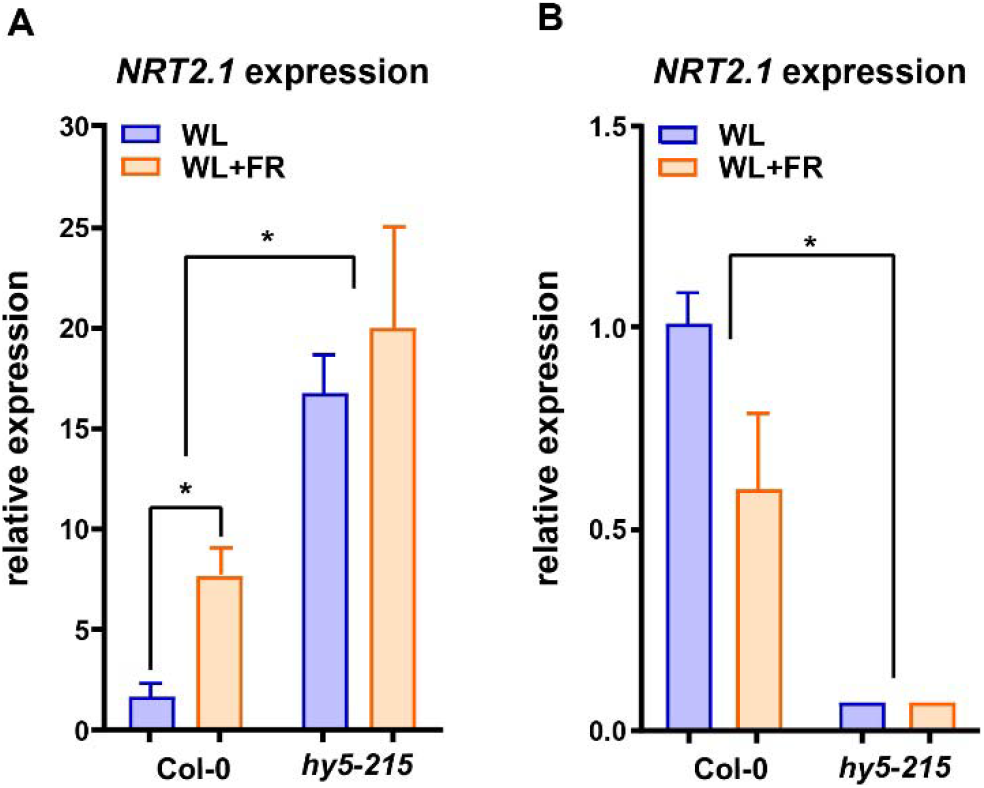
qPCR expression analysis of *NRT2.1* using RNA of 5-day old seedling roots of Col-0 and *hy5-215* treated with combined WL or WL+FR, either on normal nitrate medium (A), or on ½ MS medium (B). *p<0.05 with a one-way ANOVA plus post-hoc tukey test.

**Figure S4.**
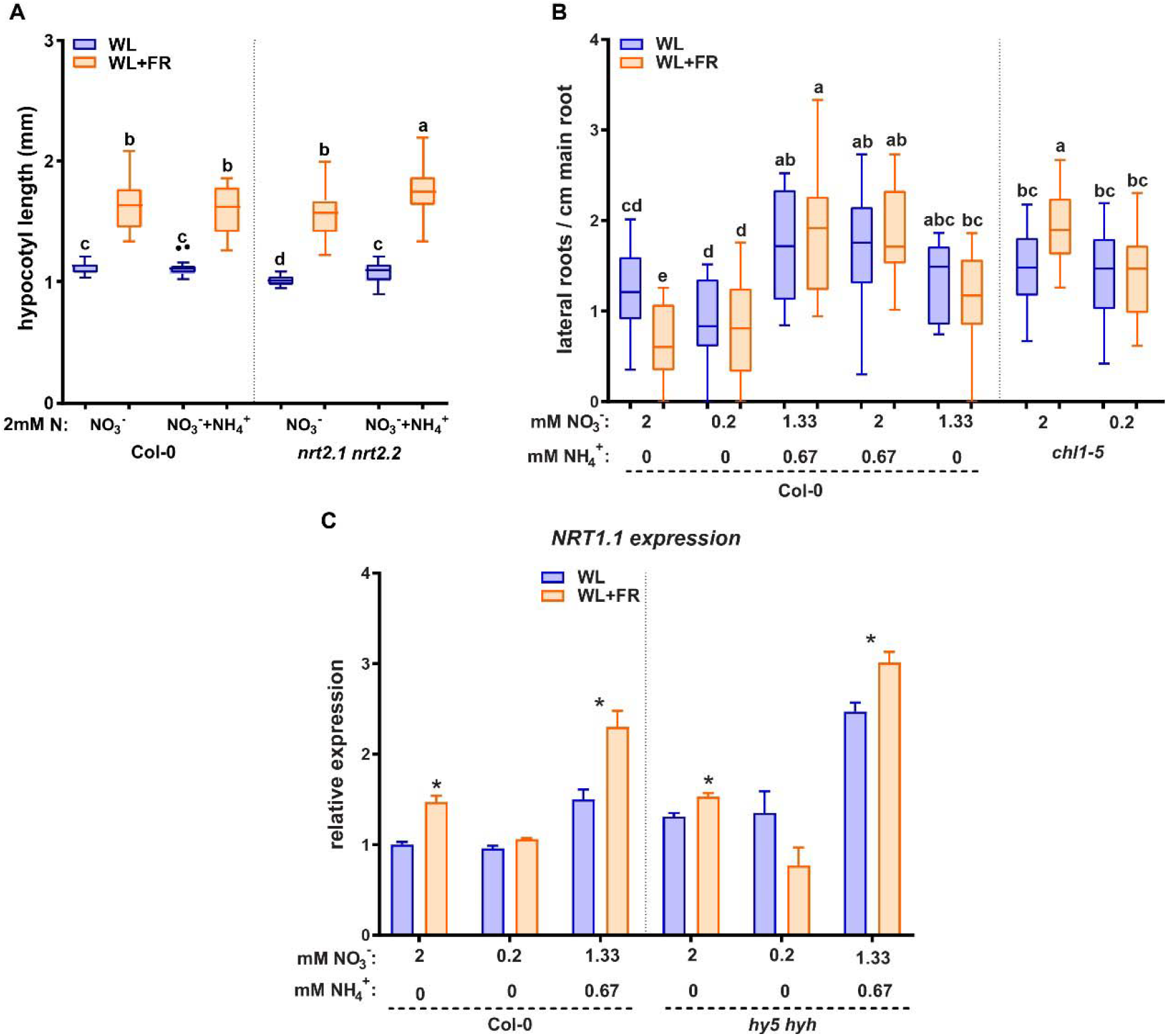
(A) Hypocotyl length of experiment shown in Figure 5. (B) Lateral root density of 8-day old wild type Col-0 seedlings grown on normal nitrate (2 mM), low (02 mM), mixed nitrate ammonium (1.33 mM + 0.67 mM NH_4_ ^+^), ammonium addition to normal nitrate (2 mM + 0.67 mM NH_4_^+^) and 1.33 mM nitrate. In the same experiment *chl1-5* (*nrt1.1*) was grown on normal (2 mM) and low (02 mM) nitrate. Letters denote statistically significant groups based on a mixed model 2-way ANOVA with posthoc tukey test (p<0.05). (C) Expression analysis of *NRT1.1* on the same material as figure 4D-E (*p<0.05 vs 2mM Col-0 WL with a one-way ANOVA plus post-hoc tukey test).

## Notes

### Competing Interest Statement

The authors have declared no competing interest.

