## supplemental figures for "Regulation of lateral root development by shoot-sensed far-red light via HY5 is nitrate-dependent and involves the NRT2.1 nitrate transporter"

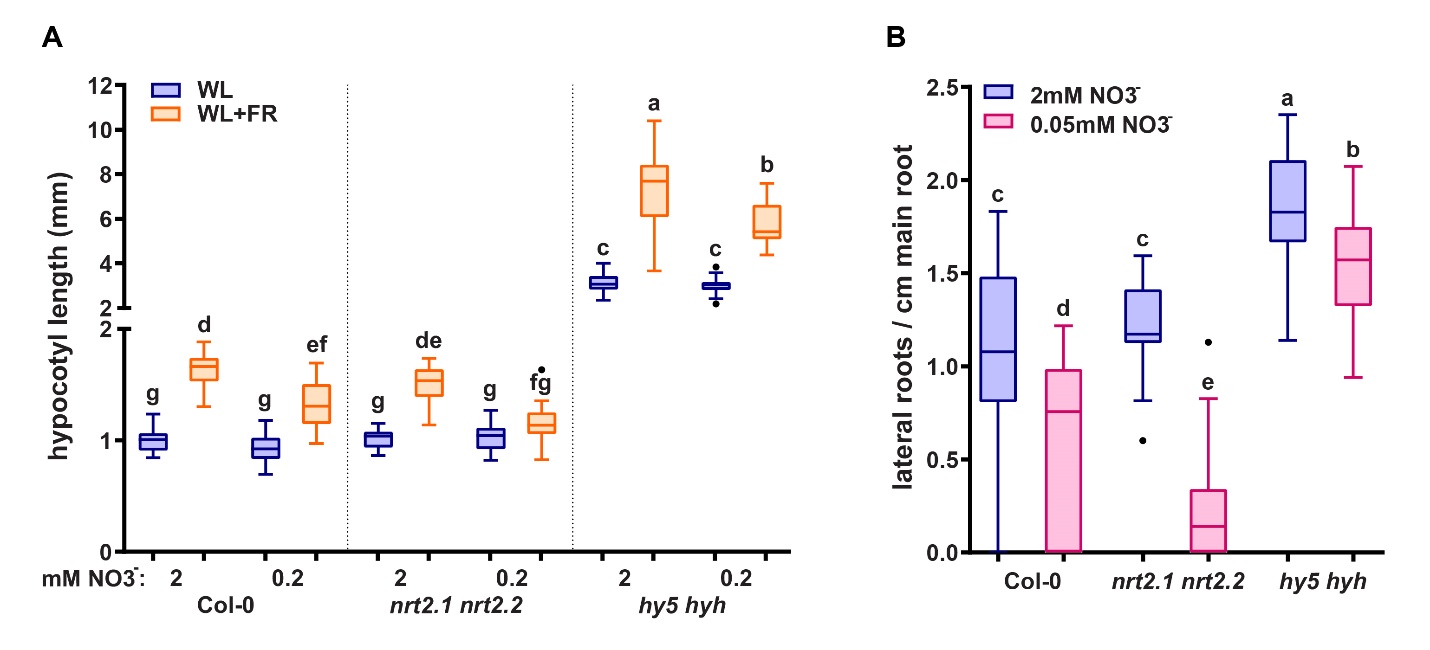

**Figure S1.** (A) Hypocotyl length of eight day old seedlings grown in WL on normal (2 mM) or low (0.2 mM) nitrate. (B) Lateral root density of eight day old seedlings grown in WL on normal (2 mM) or very low (0.05 mM) nitrate. Letters denote statistically significant groups based on a mixed model 2-way ANOVA with posthoc tukey test (p<0.05). Scale bar = 1cm.

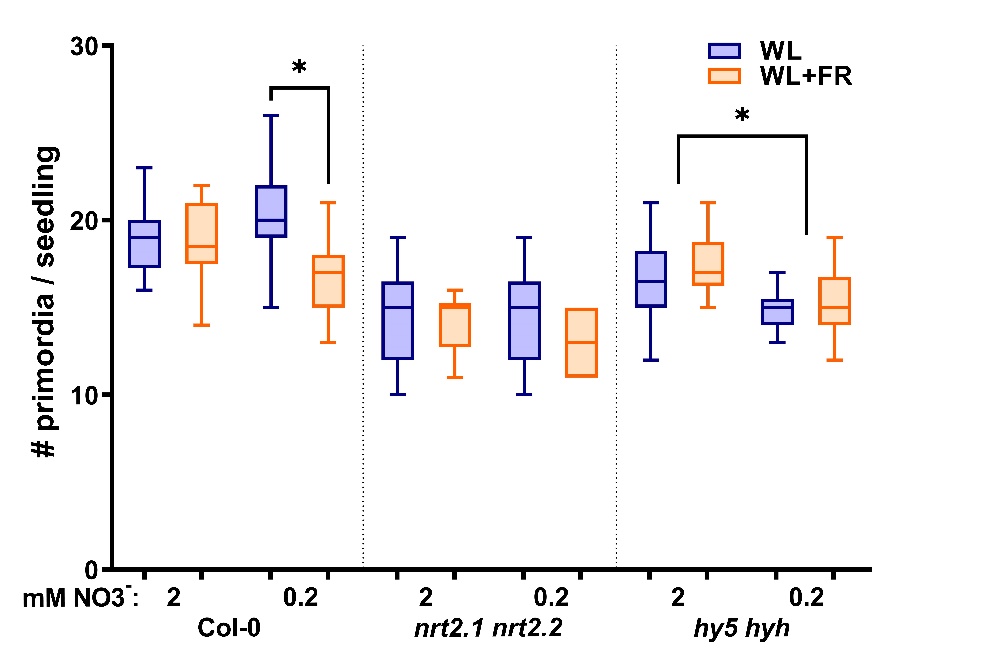

**Figure S2.** Total number of primordia per seedling from the experiment shown in figure 3. *Statistics*: mixed model 2-way ANOVA with post hoc Tukey test within the stage groups (*p<0.05).

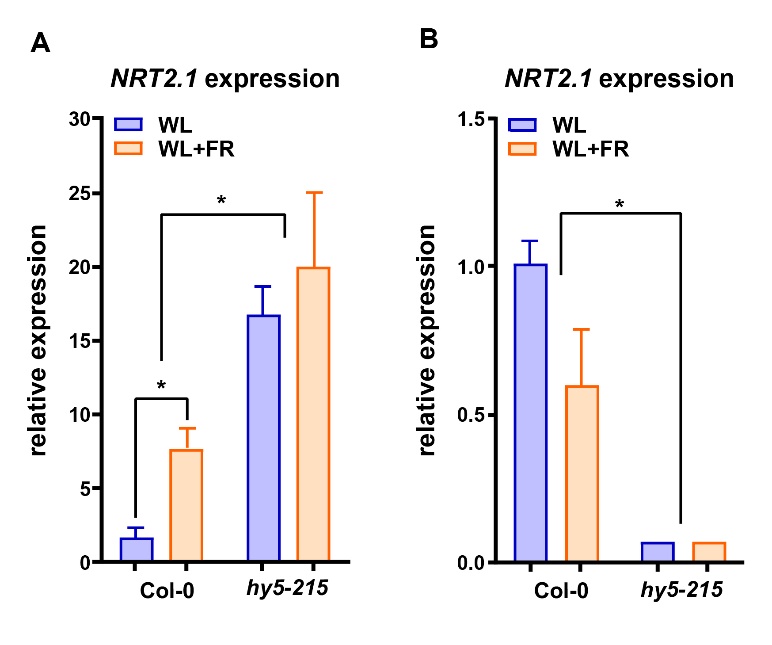

**Figure S3.** qPCR expression analysis of *NRT2.1* using RNA of 5-day old seedling roots of Col-0 and *hy5-215* treated with combined WL or WL+FR, either on normal nitrate medium (A), or on ½ MS medium (B). *p<0.05 with a one-way ANOVA plus post-hoc tukey test.

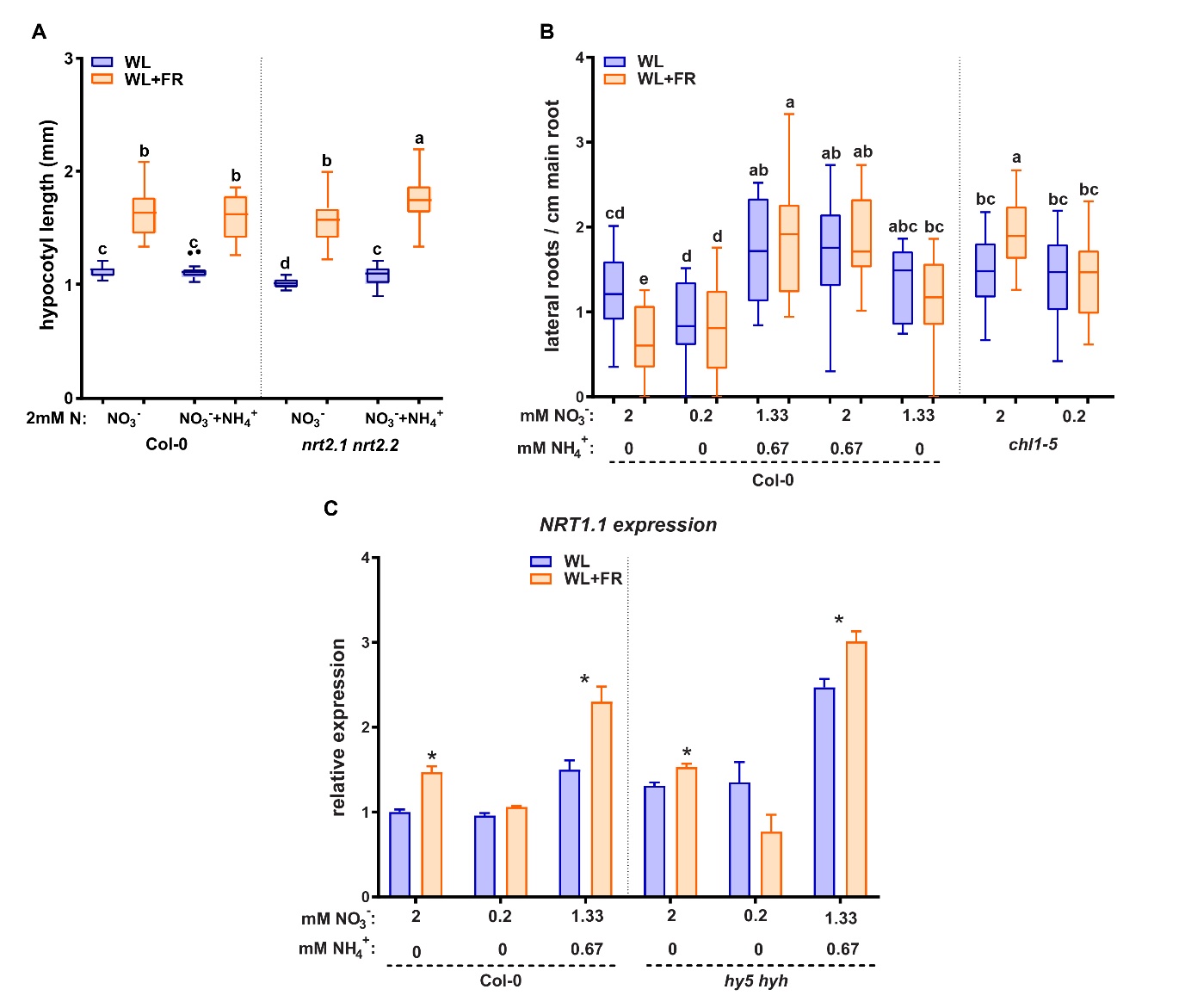

**Figure S4.** (A) Hypocotyl length of experiment shown in Figure 5. (B) Lateral root density of 8-day old wild type Col-0 seedlings grown on normal nitrate (2 mM), low (02 mM), mixed nitrate ammonium (1.33 mM + 0.67 mM NH_4_^+^), ammonium addition to normal nitrate (2 mM + 0.67 mM NH_4_^+^) and 1.33 mM nitrate. In the same experiment *chl1-5* (*nrt1.1*) was grown on normal (2 mM) and low (02 mM) nitrate. Letters denote statistically significant groups based on a mixed model 2-way ANOVA with posthoc tukey test (p<0.05). (C) Expression analysis of *NRT1.1* on the same material as figure 4D-E ( *p<0.05 vs 2mM Col-0 WL with a one-way ANOVA plus post-hoc tukey test).
